# Integrated analysis of genomics, longitudinal metabolomics, and Alzheimer’s risk factors among 1,111 cohort participants

**DOI:** 10.1101/436923

**Authors:** Burcu F. Darst, Qiongshi Lu, Sterling C. Johnson, Corinne D. Engelman

## Abstract

Although Alzheimer’s disease (AD) is highly heritable, genetic variants known to be associated with AD only explain a small proportion of its heritability. Genetic factors may only convey disease risk in individuals with certain environmental exposures, suggesting that a multi-omics approach could reveal underlying mechanisms contributing to complex traits, such as AD. We developed an integrated network to investigate relationships between metabolomics, genomics, and AD risk factors using Wisconsin Registry for Alzheimer’s Prevention participants. Analyses included 1,111 non-Hispanic Caucasian participants with whole blood expression for 11,376 genes (imputed from dense genome-wide genotyping), 1,097 fasting plasma metabolites, and 17 AD risk factors. A subset of 155 individuals also had 364 fasting cerebral spinal fluid (CSF) metabolites. After adjusting each of these 12,854 variables for potential confounders, we developed an undirected graphical network, representing all significant pairwise correlations upon adjusting for multiple testing. There were many instances of genes being indirectly linked to AD risk factors through metabolites, suggesting that genes may influence AD risk through particular metabolites. Follow-up analyses suggested that glycine mediates the relationship between *CPS1* and measures of cardiovascular and diabetes risk, including body mass index, waist-hip ratio, inflammation, and insulin resistance. Further, 38 CSF metabolites explained more than 60% of the variance of CSF levels of tau, a detrimental protein that accumulates in the brain of AD patients and is necessary for its diagnosis. These results further our understanding of underlying mechanisms contributing to AD risk while demonstrating the utility of generating and integrating multiple omics data types.

## Introduction

Genome-wide association studies (GWAS) have identified tens of thousands of single nucleotide polymorphism (SNP)-trait associations(1). However, these variants tend to have very small effect sizes and typically explain a small portion of trait heritability. Late onset Alzheimer’s disease (AD) is an example of such a trait: 53% of its phenotypic variance can be explained by genomic variants, collectively (*i.e.*, SNP heritability); yet, the 21 GWAS variants identified in a meta-analysis to be associated with AD only account for 31% of its genetic variance, leaving 69% unaccounted for(2). In order to more comprehensively understand the disease risk conveyed by genetic factors, it is crucial to consider genomics in combination with other omics data types and to use integrative multi-omics approaches that can capture intricate relationships.

Although there has been great interest recently in the integration of multi-omics datasets, progress in this field is still fairly limited and it faces many challenges(3–8). However, studies have been able to show that the use of multiple omics data types is more predictive than single data types(5, 9). A recent study with dense longitudinal omics data displayed the utility of integrating such data with regards to personalized medicine(10). Although limited by its sample size of 108 participants, this investigation identified meaningful systems biology relationships that were able to improve the health of its participants. As it is becoming more feasible and common to acquire multiple omics data types, it is essential that we move towards systems biology approaches of understanding complex diseases, rather than focusing on single data types that are unable to capture the intricacies imposed by biology.

Recent technological advances have made metabolomics studies increasingly favorable among investigations of AD(11), obesity(12), and cardiovascular disease(13), to name a few. An appeal of the metabolome is that of the biological systems, metabolomics could offer an effective way to accurately capture individual-level environmental exposures; it is the most proximal to the development of the phenotype(14) and many metabolites have a low heritability(15, 16), implying that such metabolites are more strongly influenced by the environment than genomics. Metabolomic variations that precede disease onset could prove to be highly informative for predictive models as well as preventative and therapeutic medicine. Pathological changes that cause AD are known to begin decades before the diagnosis of AD(17). As such, an integrated approach of studying the genomics and metabolomics of risk factors that precede an AD diagnosis could provide a better understanding of the underlying biological and environmental mechanisms that lead to the onset of AD.

We developed an integrative network to investigate relationships between plasma metabolomics, cerebral spinal fluid (CSF) metabolomics, genomics, and AD risk factors using 1,111 participants with deep longitudinal phenotypes from the Wisconsin Registry for Alzheimer’s Prevention (WRAP). AD risk factors included neuropsychological measures of cognitive function, CSF levels of the two proteins required for an AD diagnosis that are known to accumulate in the brains of AD patients, amyloid-beta (Aβ) and tau, and measures of cardiovascular disease and diabetes risk, two diseases that are known to increase AD risk. Further, in order to understand whether plasma metabolite levels are representative of metabolites in CSF, which may be a more relevant tissue for neurological diseases, we also assessed the correlation of plasma and CSF metabolite levels.

## Materials and Methods

### Participants

Study participants were from WRAP, a longitudinal study of initially dementia free middle-aged adults that allows for the enrollment of siblings and is enriched for a parental history of Alzheimer’s disease. Further details of the study design and methods used have been previously described(18, 19). Participants included in this analysis had genetic ancestry that was primarily of European descent, had both genomic and metabolomic data available, and up to seventeen AD risk factors (Table 1; of note, cholesterol is not included in this table because it was measured on the metabolite panel). This study was conducted with the approval of the University of Wisconsin Institutional Review Board, and all subjects provided signed informed consent before participation.

**Table 1.**
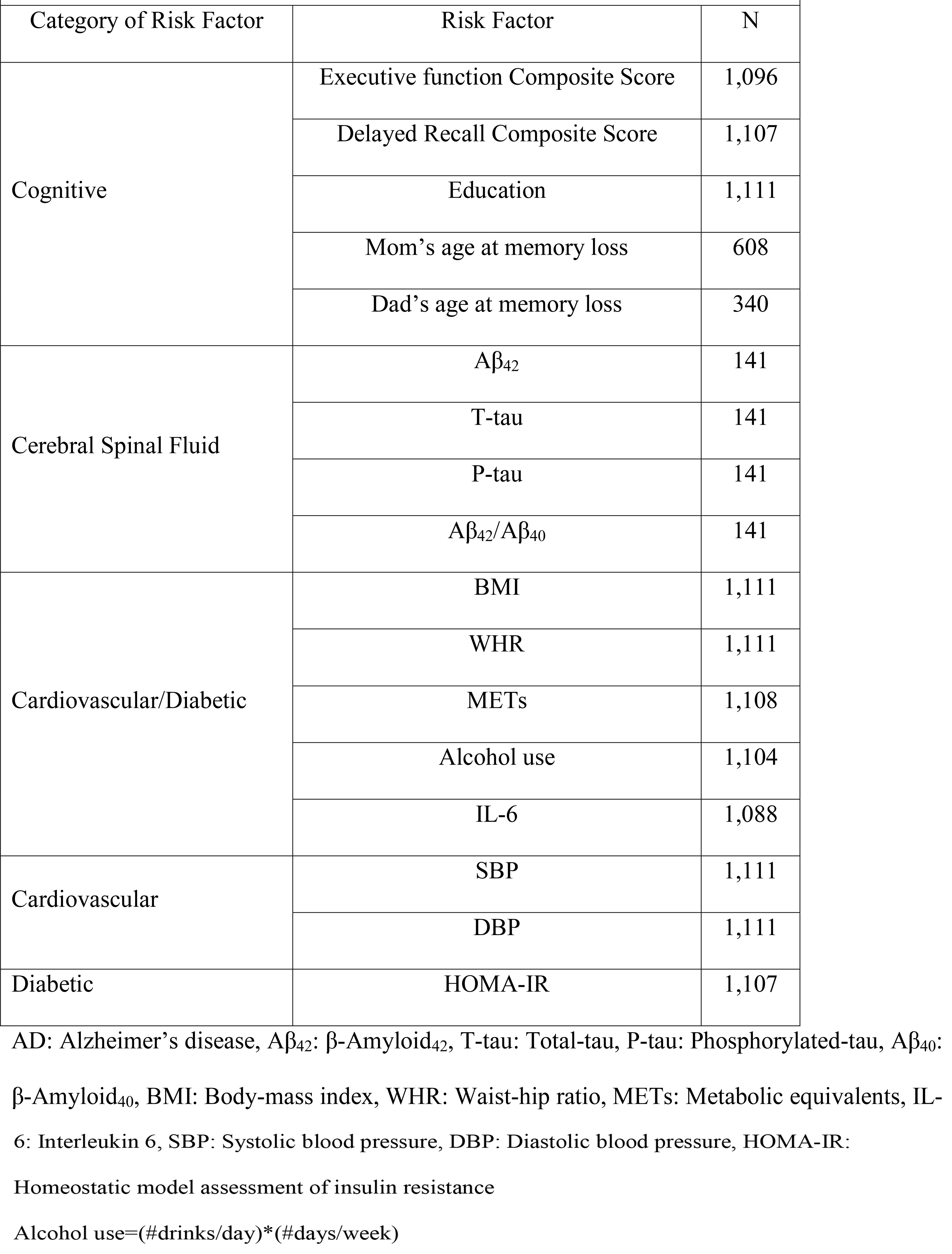
Seventeen AD Risk Factors Included in Network Analysis

| Table 1. Seventeen AD Risk Factors Included in Network Analysis |  |  |
| --- | --- | --- |
| Category of Risk Factor | Risk Factor | N |
| Cognitive | Executive function Composite Score | 1,096 |
|  | Delayed Recall Composite Score | 1,107 |
|  | Education | 1,111 |
|  | Mom's age at memory loss | 608 |
|  | Dad's age at memory loss | 340 |
| Cerebral Spinal Fluid | Aβ<sub>42</sub> | 141 |
|  | T-tau | 141 |
|  | P-tau | 141 |
| | Aβ<sub>42</sub>/Aβ<sub>40</sub> | 141 |
| Cardiovascular/Diabetic | BMI | 1,111 |
|  | WHR | 1,111 |
|  | METs | 1,108 |
|  | Alcohol use | 1,104 |
|  | IL-6 | 1,088 |
| Cardiovascular | SBP | 1,111 |
|  | DBP | 1,111 |
| Diabetic | HOMA-IR | 1,107 |

AD: Alzheimer's disease, Aβ <sub>42</sub>: $\beta$ -Amyloid<sub>42</sub>, T-tau: Total-tau, P-tau: Phosphorylated-tau, Aβ <sub>40</sub>:
$\beta$ -Amyloid<sub>40</sub>, BMI: Body-mass index, WHR: Waist-hip ratio, METs: Metabolic equivalents, IL-
6: Interleukin 6, SBP: Systolic blood pressure, DBP: Diastolic blood pressure, HOMA-IR:
Homeostatic model assessment of insulin resistance
Alcohol use=(#drinks/day)\*(#days/week)

### Plasma and CSF collection and sample handling

Fasting blood samples for this study were drawn the morning of each study visit. Plasma samples were stored in ethylenediaminetetraacetic acid (EDTA) tubes at −80°C. Blood was collected in 10 mL ethylenediaminetetraacetic acid (EDTA) vacutainer tubes. They were immediately placed on ice, and then centrifuged at 3000 revolutions per minute for 15 minutes at room temperature. Plasma was pipetted off within one hour of collection. Plasma samples were aliquoted into 1.0 mL polypropylene cryvolials and placed in −80°C freezers within 30 minutes of separation.

As previously described(20), CSF was collected via lumbar puncture (LP) in the morning after a 12-hour fast, not necessarily on the same day as a study visit (LPs were drawn within a median of 120 days of the study visit, ranging from 0-661 days). LPs were performed using a Sprotte 25- or 24-gauge spinal needle at the L3/4 or L4/5 interspace using gentle extraction into polypropylene syringes. CSF (22 mL) was then gently mixed and centrifuged at 2000g for 10 minutes. Supernatants were frozen in 0.5 mL aliquots in polypropylene tubes and stored at −80°C.

Plasma and CSF samples were never thawed before being shipped overnight on dry ice to Metabolon (Durham, NC), where they were again stored in −80°C freezers and thawed once before testing.

### CSF biomarker quantification

CSF Aβ_42_, total tau (T-tau), and phosphorylated tau (P-tau) were quantified with sandwich ELISAs (INNOTEST β-amyloid1-42, hTAU-Ag, and Phospho-Tau[181P], respectively; Fujirebio Europe, Ghent, Belgium). CSF levels of Aβ_42_ and Aβ_40_ (a less amyloidogenic Aβ fragment as compared to Aβ_42_) were used to calculate the ratio of Aβ_42_/Aβ_40_ were quantified by electrochemiluminescence (ECL) using an Aβ triplex assay (MSD Human Aβ peptide Ultra-Sensitive Kit, Meso Scale Discovery, Gaithersburg, MD). A total of 223 samples with CSF biomarkers among 141 individuals were available for this analysis.

### Plasma and CSF metabolomic profiling and quality control

Untargeted plasma and CSF metabolomic analyses and quantification were performed by Metabolon (Durham, NC) using Ultrahigh Performance Liquid Chromatography-Tandom Mass Spectrometry (UPLC-MS/MS)(21); details are outlined in the Supplemental Note. Metabolites within eight super pathways were identified: amino acids, carbohydrates, cofactors and vitamins, energy, lipids, nucleotides, peptides, and xenobiotics.

Up to three longitudinal plasma samples were available for each participant. Plasma metabolites with an interquartile range of zero (*i.e.*, those with very low or no variability) were excluded from analyses (178 metabolites). After removing these metabolites, samples were missing a median of 11.7% plasma metabolites, while plasma metabolites were missing in a median of 1.2% of samples.

Up to four longitudinal CSF samples were available for each participant. Similarly, CSF metabolites with an interquartile range of zero were excluded from analyses (48 CSF metabolites). After removing these metabolites, samples were missing a median of 6.9% CSF metabolites, while CSF metabolites were missing in a median of 0.3% of samples.

Missing plasma and CSF metabolite values were imputed to the lowest level of detection for each metabolite(22). Metabolite values were median-scaled and log-transformed to normalize metabolite distributions(23). If a participant reported that they did not fast or withhold medications and caffeine for at least eight hours prior to the blood draw, the plasma sample was excluded from analyses (159 plasma samples), leaving 1,097 plasma metabolites among 2,189 plasma samples (1,111 individuals) for analyses. Similarly, if a participant reported that they did not fast for at least eight hours prior to the LP, the CSF sample was excluded from analyses (4 CSF samples), leaving 364 CSF metabolites among 346 CSF samples (155 individuals) for analyses.

### CSF and plasma metabolite correlations

A total of 326 metabolites were captured in both CSF and plasma. The correlations of these metabolites between tissue types were calculated using the Pearson correlation coefficient. In order to reduce variability due to the time interval between plasma and CSF sample collection, correlations were based on 141 pairs of plasma and CSF samples that were collected within a timespan of four months of each other. After removing these samples, plasma and CSF samples were collected a median of 27 days apart.

### DNA collection and genomics quality control

DNA was extracted from whole blood samples using the PUREGENE^®^ DNA Isolation Kit (Gentra Systems, Inc., Minneapolis, MN). DNA concentrations were quantified using the Invitrogen™ Quant-iT™ PicoGreen™ dsDNA Assay Kit (Thermo Fisher Scientific, Hampton, NH) analyzed on the Synergy 2 Multi-Detection Microplate Reader (Biotek Instruments, Winooski, VT). Samples were normalized to 50 ng/ul following quantification.

A total of 1,340 samples were genotyped using the Illumina Multi-Ethnic Genotyping Array at the University of Wisconsin Biotechnology Center (Figure S1). Thirty-six blinded duplicate samples were used to calculate a concordance rate of 99.99%, and discordant genotypes were set to missing. Sixteen samples missing >5% of variants were excluded, while 35,105 variants missing in >5% of individuals were excluded. No samples were removed due to outlying heterozygosity. Six samples were excluded due to inconsistencies between self-reported and genetic sex.

Due to the sibling relationships present in the WRAP cohort, genetic ancestry was assessed using Principal Components Analysis in Related Samples (PC-AiR), a method that makes robust inferences about population structure in the presence of relatedness(24). This approach included several iterative steps and was based on 63,503 linkage disequilibrium (LD) pruned (r^2^<0.10) and common (MAF>0.05) variants, using the 1000 Genomes data as reference populations(25). First, kinship coefficients (KCs) were calculated between all pairs of individuals using genomic data with the Kinship-based Inference for Gwas (KING)-robust method(26). PC-AiR was used to perform principal components analysis (PCA) on the reference populations along with a subset of unrelated individuals identified by the KCs. Resulting principal components (PCs) were used to project PC values onto the remaining related individuals. All PCs were then used to recalculate the KCs taking ancestry into account using the PC-Relate method, which estimates KCs robust to population structure(27). PCA was performed again using the updated KCs, and KCs were also estimated again using updated PCs. The resulting PCs identified 1,198 WRAP participants whose genetic ancestry was primarily of European descent. This procedure was repeated within this subset of participants (excluding 1000 Genomes individuals) to obtain PC estimates used to adjust for population stratification in subsequent genomic analyses. Among European descendants, 160 variants were not in Hardy-Weinberg equilibrium (HWE) and 327,064 were monomorphic and thus, removed.

A total of 1,294,660 bi-allelic autosomal variants among 1,198 European descendants remained for imputation, which was performed with the Michigan Imputation Server v1.0.3(28), using the Haplotype Reference Consortium (HRC) v. r1.1 2016(29) as the reference panel and Eagle2 v2.3(30) for phasing. Prior to imputation, the HRC Imputation Checking Tool(31) was used to identify variants that did not match those in HRC, were palindromic, differed in MAF>0.20, or that had non-matching alleles when compared to the same variant in HRC, leaving 898,220 for imputation. A total of 39,131,578 variants were imputed. Variants with a quality score R^2^<0.80, MAF<0.001, or that were out of HWE were excluded, leaving 10,400,394 imputed variants. These were combined with the genotyped variants, leading to 10,499,994 imputed and genotyped variants for analyses. Data cleaning and file preparation were completed using PLINK v1.9(32) and VCFtools v0.1.14(33). Coordinates are based on GRCh37 assembly hg19.

### Whole blood gene expression imputation

The resulting 10,499,994 imputed and genotyped variants were used to impute whole blood gene expression using PrediXcan(34) with the Depression Genes and Networks reference dataset(35), PrediXcan’s largest reference sample consisting of 922 individuals with RNA sequencing on whole blood and GWAS data. PrediXcan filters results to only include genes that are imputed with reasonable accuracy, using a false discovery rate of 0.05. After removing genes with zero variability between individuals (162 genes), whole blood gene expression data for 11,376 genes were available for analyses.

### Integrative network analysis

The analytic approach we used for our network analysis was similar to that of Price et al., 2017(10). A total of 12,856 variables, including 11,376 expressed genes, 1,097 plasma metabolites, 364 CSF metabolites, and 17 AD risk factors, were available for the network analysis. Linear mixed models, as implemented by the lme4 package in R(36), were used to adjust each variable for age and sex and included a random intercept for individual to account for repeated measures and family to account for sibling relationships. Further adjustments were made specific to the variable being assessed: imputed gene expression was also adjusted for the first four principal components to account for ancestry; CSF and plasma metabolites were adjusted for cholesterol lowering medication use and sample storage time; the executive function and delayed recall composite scores were adjusted for practice effects; and systolic and diastolic blood pressure were adjusted for ace inhibitor and beta blocker medication use. For longitudinal traits (such as metabolites), random intercepts were used as the new outcomes for each individual, whereas for constant traits (such as imputed gene expression values), residuals were used as the new outcomes for each individual. These adjusted outcomes were used to assess all 82,606,231 pairwise correlations between traits using Spearman rank, and significance was determined using a Bonferroni-adjusted *P*-value (0.05/82,606,231=6.1e-10). To identify relationships between omics data, significant inter-omic associations and significant associations with an AD risk factor were used to develop an integrative network, which was created using the igraph R package(37). Dense subgraphs were identified using a community detection algorithm that maximizes the modularity of the network, such that there is high connectivity within communities (or groups of distinct variables), but low connectivity between communities(38).

### Targeted mediation and interaction analyses

Results from the integrated network analysis were used to identify potential mediation and interactions between imputed gene expression and metabolite levels that could impact AD risk factors, as a proof of concept. Although our network analysis suggested many potentially meaningful mediation or interaction relationships, we only investigated gene-metabolite correlations with the most consistent biological support from the GWAS catalog(1) (www.ebi.ac.uk/gwas, date accessed: May 9, 2018), to illustrate the utility of the network analysis results. Such relationships were investigated using the longitudinal data (2,198 observations among 1,111 individuals) with linear mixed models, again as implemented by the lme4 package in R(36), including random intercepts for within-individual repeated measures and within-family relationships. To assess whether a metabolite mediated the relationship between imputed gene expression and an AD risk factor, models were run to assess whether: 1) the gene predicted the AD risk factor, 2) the gene predicted metabolite levels, 3) the metabolite predicted the AD risk factor, and 4) the gene predicted the AD risk factor while adjusting for the metabolite. The causal mediation effect, or the indirect effect of a gene on an AD risk factor through a metabolite, was calculated as the difference between the effect of the gene in model 1 and model 4, as implemented in the R mediation package(39). To determine whether this difference was significant, standard errors and *P*-values were estimated using the quasi-Bayesian Monte Carlo method with 1,000 simulations. Because the mediation package can only handle mixed models with one random effect, the mediation analysis was run with models 1 and 4 excluding the random effect for family. As a sensitivity analysis, the mediation analysis was rerun limiting models 1 and 4 to unrelated individuals (n=898 with 1,774 observations). A fifth linear mixed model was used to assess interactions by adding a gene*metabolite interaction term to model 4. Model 5 did not use the mediation package and was thus able to include random intercepts for both within-individual repeated measures and within-family relationships. All models including a gene had covariates for age, sex, and the first four PCs, while models including a metabolite had covariates for age, sex, cholesterol lowering medication use, and sample storage time.

## Results

### Participants

A total of 1,111 WRAP participants had both genomic and plasma metabolomic data. At baseline, 68.9% of participants were female and participants were 61.0 years old with a bachelor’s degree, on average (Table 2). Participants each had 1,097 plasma metabolites available for analyses, 347 (31.6%) of which were of unknown chemical structure, whole blood gene expression for 11,376 genes, and up to 17 AD risk factors. A subset of 155 individuals also had 364 CSF metabolites available for analyses, 56 (15.4%) of which were of unknown chemical structure. Participants with CSF metabolomic data had similar characteristics as the full sample (Table 2). Properties of each plasma and CSF metabolite, such as biochemical name, super pathway, and sub pathway are described in Table S1, and numbers of metabolites within each super pathway are summarized in Table S2.

**Table 2.**
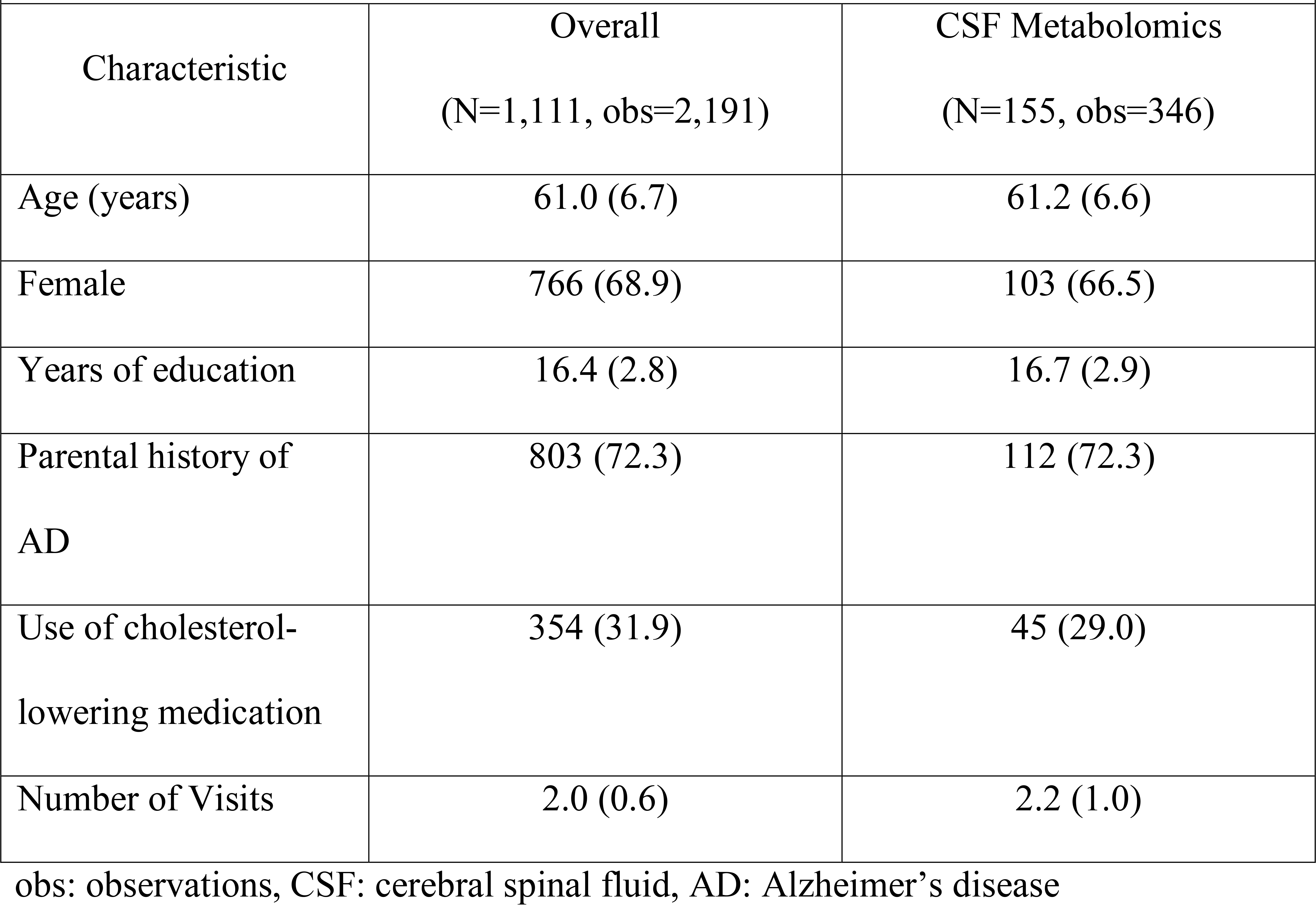
WRAP Participant Characteristics at Baseline Sample. Mean (SD) or N (%).

### Correlation between plasma and CSF metabolomics

The median correlation between the 326 metabolites common to both plasma and CSF was r=0.26, with some variability existing between different metabolite pathways (Figure 1). Xenobiotics had the highest median correlation (r=0.53), while lipids had the lowest (r=0.11). Overall, metabolite correlations ranged from |r|=0.0002 (inosine, a nucleotide) to |r|=0.88 (quinate, a xenobiotic). Interestingly, one of the highest correlations was caffeine (r=0.81).

**Figure 1.**
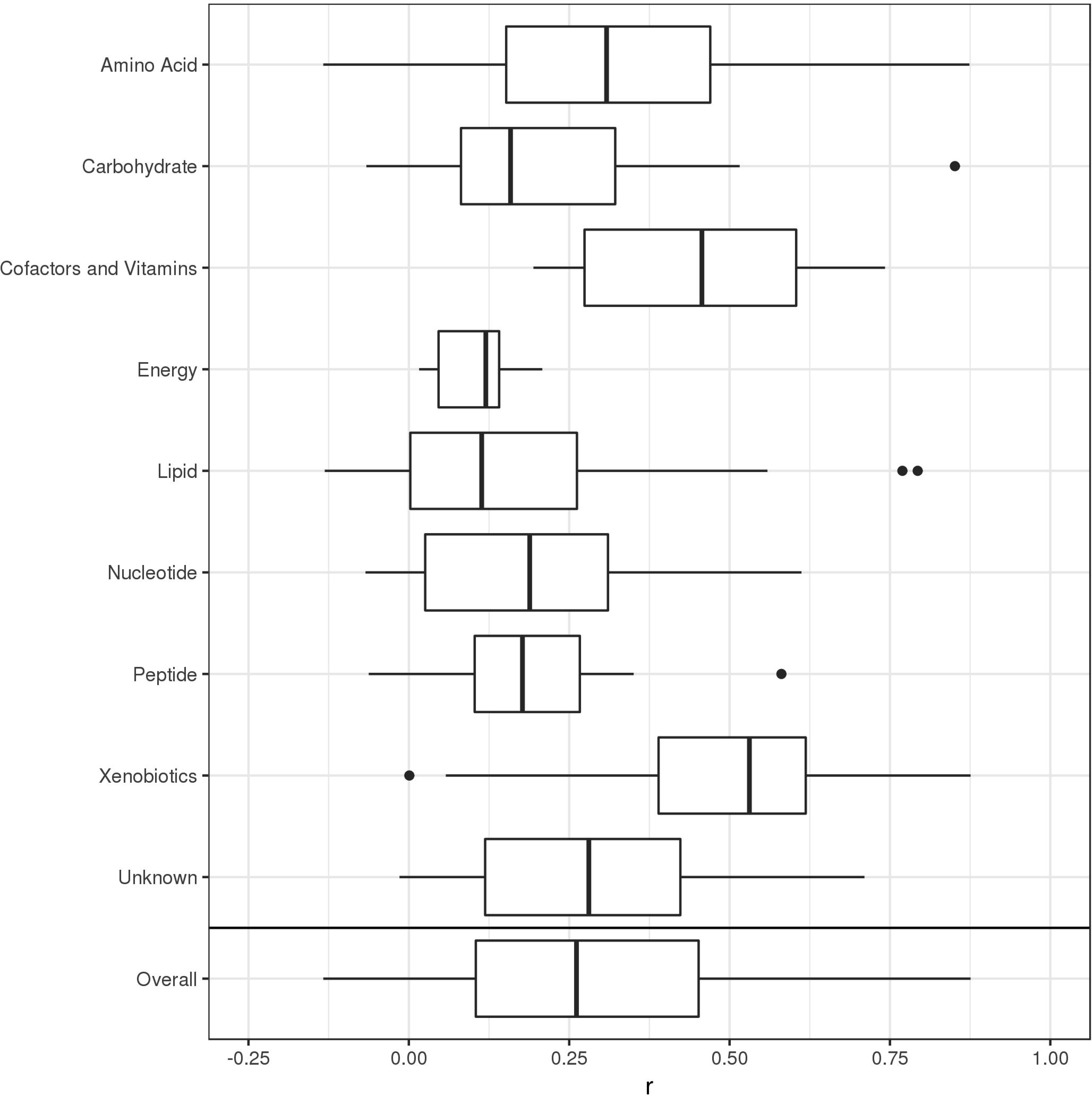
Correlations between plasma and CSF metabolites by super pathway. Vertical bars represent median correlations; box width represents the first and third quartiles; horizontal bars (whiskers) represent the range of correlations that are within 1.5 times the interquartile range; and dots represent outlier correlations that exceed 1.5 times the interquartile range.

Correlations between each of the 326 CSF and plasma metabolites are described in Table S3.

### Integrated network

After applying a Bonferroni correction for multiple testing, a total of 90,308 significant correlations (edges) among 10,869 variables (nodes) were used to develop an overall ‘hairball’ network (Figure S2). Notably, although there were far fewer metabolites than genes in the network (1,387 metabolites versus 9,481 genes), there were more edges between metabolites than genes (49,499 versus 37,473 edges, respectively).

The inter-omic network is shown in Figure 2 (a labeled version is shown in Figure S3), and its corresponding community partitions are shown in Figure S4. This network had 1,224 edges and 635 nodes, including 171 metabolite-gene and 833 metabolite-AD risk factor edges. Of these, there were only four CSF metabolite-gene edges and 73 CSF metabolite-AD risk factor edges, likely due to the much smaller number of CSF metabolomic samples. No genes were directly linked to AD risk factors; however, many genes were indirectly linked to AD risk factors through metabolites, as described below. Each of the 1,224 correlations is described in Table S4.

**Figure 2.**
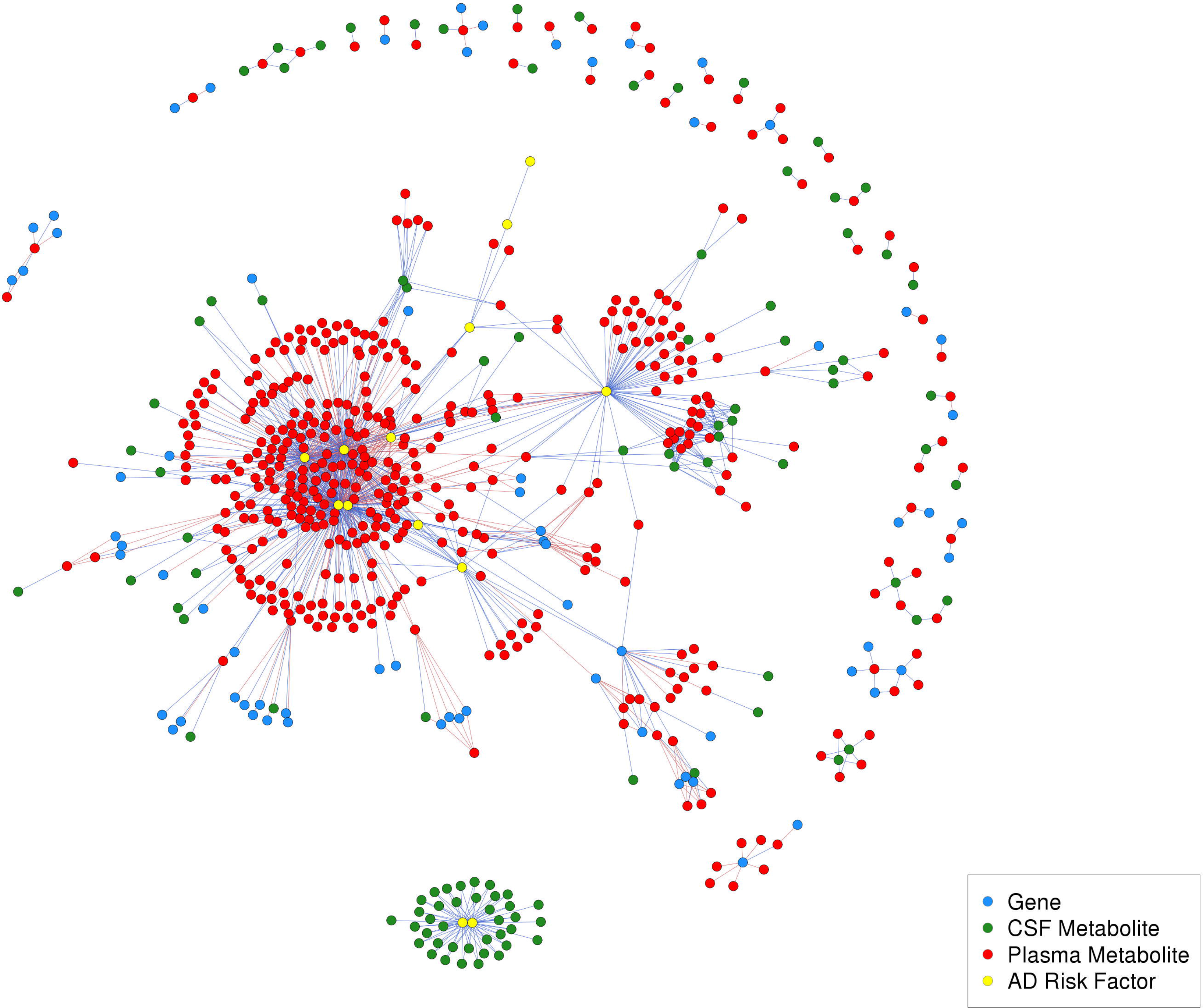
Inter-omic network. This network has 1,224 edges and 635 nodes, which included 171 metabolite-gene edges, 833 metabolite-AD risk factor edges. Of these, 73 were CSF metabolite-AD risk factor edges (CSF T-tau and P-tau, exclusively) and 4 were CSF metabolite-gene edges. Red edges indicate negative correlations and blue edges indicate positive correlations.

The largest community contained 680 edges among 289 nodes, which included 264 plasma metabolites, ten CSF metabolites, eight genes, and seven AD risk factors related to cardiovascular disease and diabetes: body mass index (BMI), waist-hip ratio (WHR), homeostatic model assessment of insulin resistance (HOMA-IR), interleukin 6 (IL-6), metabolic equivalents (METs), diastolic blood pressure (DBP), and systolic blood pressure (SBP) (Figure S5). Expression levels of these eight genes were all indirectly linked to AD risk factors within this community through plasma metabolites. *CPS1* expression levels were negatively correlated with plasma gamma-glutamylglycine, proprionylglycine, and glycine levels, all of which were negatively correlated with BMI, WHR, IL-6, and/or HOMA-IR (Figure 3). *TMEM229B* and *PLEKHH1* were both negatively correlated with two glycerophosphatidylcholines (1-(1-enyl-palmitoyl)-2-palmitoleoyl-GPC (P-16:0/16:1) and 1-(1-enyl-palmitoyl)-2-palmitoyl-GPC (P-16:0/16:0)), which were also negatively correlated with BMI, WHR, and/or HOMA-IR. *NAALAD2* was negatively correlated with an amino acid beta-citrylglutamate, which was positively correlated with BMI, WHR, IL-6, and HOMA-IR. *ZNF655* and *ZKSCAN1* were both positively correlated with X-12063, which was also positively correlated with BMI, WHR, and HOMA-IR. *CHRNA5* was positively correlated with 5-hydroxylysine, which was positively correlated with BMI, WHR, IL-6, and HOMA-IR, and negatively correlated with METs. *ARVCF* was negatively correlated with X-11593, which was positively correlated with BMI, IL-6, and HOMA-IR.

**Figure 3.**
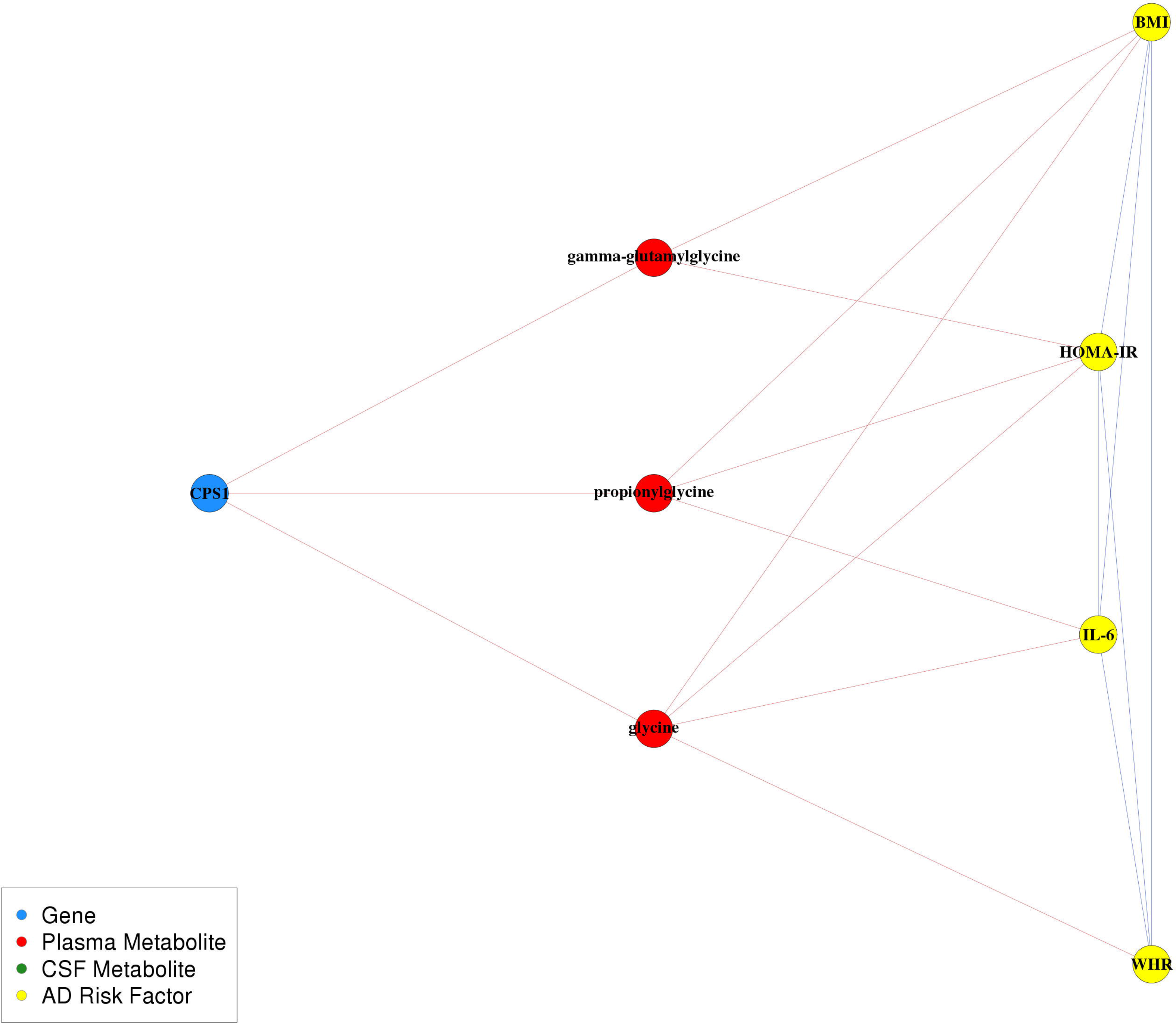
*CPS1*, glycine, and cardiovascular and diabetes sub-network. Relationships within this pathway are highly cited; however, the pathway as a whole is not understood as well. Red edges indicate negative correlations and blue edges indicate positive correlations.

Several genes outside of the cardiovascular and diabetes community were indirectly linked to AD risk factors within this community. Gene expression of *FOSL2*, *KRTCAP3*, and *ZNF513* were positively correlated, while *IFT172*, *NRBP1*, *PPM1G*, and *ZNF512* were negatively correlated, with levels of plasma mannose, a carbohydrate that was positively correlated with BMI, WHR, IL-6, and HOMA-IR (Figure S6A). *CABP1*, *SPPL3*, and *UNC119B* expression levels were negatively correlated with plasma butyrylcarnitine (C4), which was positively correlated with BMI, WHR, IL-6, and HOMA-IR (Figure S6B). *SLC27A4*, *PHYHD1*, *ENDOG*, and *SH3GLB2* expression levels were negatively correlated with plasma 2’-O-methyluridine and 2’-O-methylcytidine levels, both nucleotides involved in pyrimidine metabolism, and the latter nucleotide is also negatively correlated with BMI and WHR (Figure S6C). *PHYHD1* was also negatively correlated with CSF levels of 2’-O-methylcytidine.

The only correlations identified among the CSF biomarkers (*i.e.*, amyloid and tau) are shown in Figure 4. Higher CSF T-tau and P-tau levels were correlated with higher levels of 38 CSF metabolites, collectively. These metabolites included 13 lipids (six phosphatidylcholines, two lysophosphatidylcholines, five sphingolipids, and cholesterol), seven amino acids, five carbohydrates, one nucleotide, one energy metabolite, one cofactor and vitamin metabolite, one xenobiotic, and nine unknown metabolites. However, none of the CSF amyloid biomarkers were correlated with CSF metabolites. We investigated how much of the variance of T-tau and P-tau could be explained by these metabolites with linear mixed models that included random intercepts for within-subject repeated measures and within-family relationships, using the R^2^ statistic for mixed models as defined by Edwards et al., 2008(40) and implemented in the r2glmm R package. After removing the variation explained by age and sex, the 37 metabolites correlated with T-tau explained 60.7% of the variation of T-tau, while the 35 metabolites correlated with P-tau explained 64.0% of the variation of P-tau.

**Figure 4.**
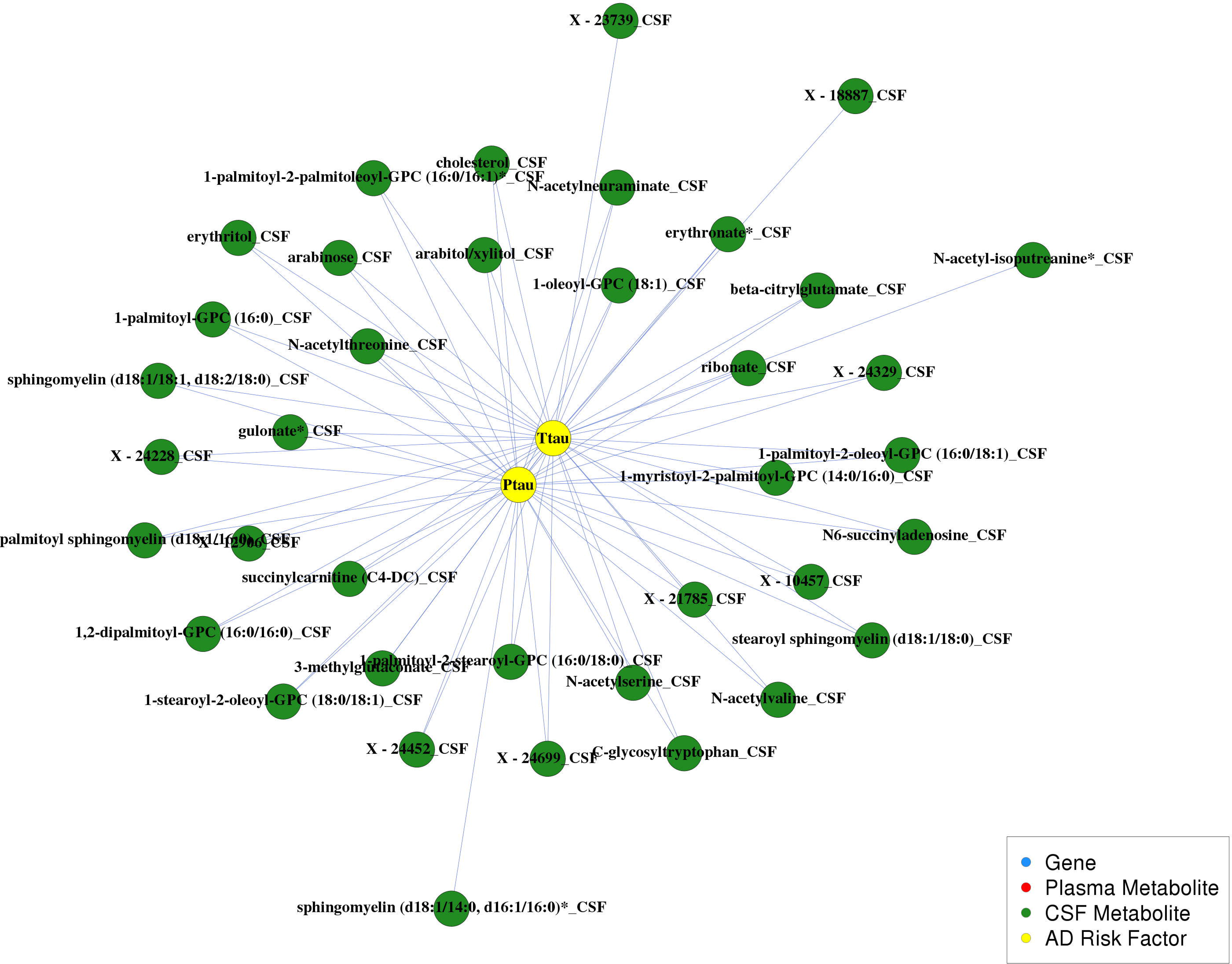
CSF biomarker community. This network has 73 edges among 38 CSF metabolites and CSF biomarkers T-tau and P-tau. Red edges indicate negative correlations and blue edges indicate positive correlations.

### Targeted mediation and interaction analyses

Targeted mediation and interaction analyses were focused on a particular pathway identified within the large cardiovascular and diabetes community involving *CPS1*, glycine plasma metabolites (glycine, proprionylglycine, and gamma-glutamylglycine), BMI, WHR, IL-6, and HOMA-IR. Associations between *CPS1* variants and glycine have been reported in at least nine studies(15, 16, 41–47), more than any of the other gene-metabolite associations identified in our network analysis, and these studies were based not only on Caucasian populations, but also on Japanese and African American populations. Many previous studies have also reported associations between glycine and cardiovascular risk factors, including BMI, waist circumference, inflammation, and HOMA-IR(45, 48–55). This evidence made this pathway a strong candidate for mediation and interaction analyses.

Figure 5 shows results from the mediation analyses using glycine as the mediator, including the total effect (*i.e.*, the effect of *CPS1* in the model unadjusted for glycine), the direct effect (*i.e.*, the effect of *CPS1* in the model adjusted for glycine), and the indirect effect (*i.e.*, the effect of *CPS1* due to the effect of *CPS1* on glycine) for BMI (Figure 5A and Figure5B), WHR (Figure 5C and Figure 5D), IL-6 (Figure 5E and Figure 5F), and HOMA-IR (Figure 5G and Figure 5H). The total effect of *CPS1* was null for each of these three outcomes, likely due to the negative association between *CPS1* and glycine coupled with the negative association between glycine and the AD risk factor, resulting in direct and indirect effects that had opposing directions(56). Our results show that lower levels of *CPS1* expression lead to increased glycine levels, and higher glycine levels lead to decreased BMI, WHR, IL-6, and HOMA-IR. Thus, with glycine as a mediator, lower levels of *CPS1* lead to decreased BMI, WHR, IL-6, and HOMA-IR. Mediation analyses using propionylglycine and gamma-glutamylglycine as the mediator showed similar results and can be found in Figure S7 and Figure S8. We did not identify any interactions between *CPS1* and the three glycine metabolites that were associated with BMI, WHR, IL-6, or HOMA-IR (all *P*-values>0.07).

**Figure 5.**
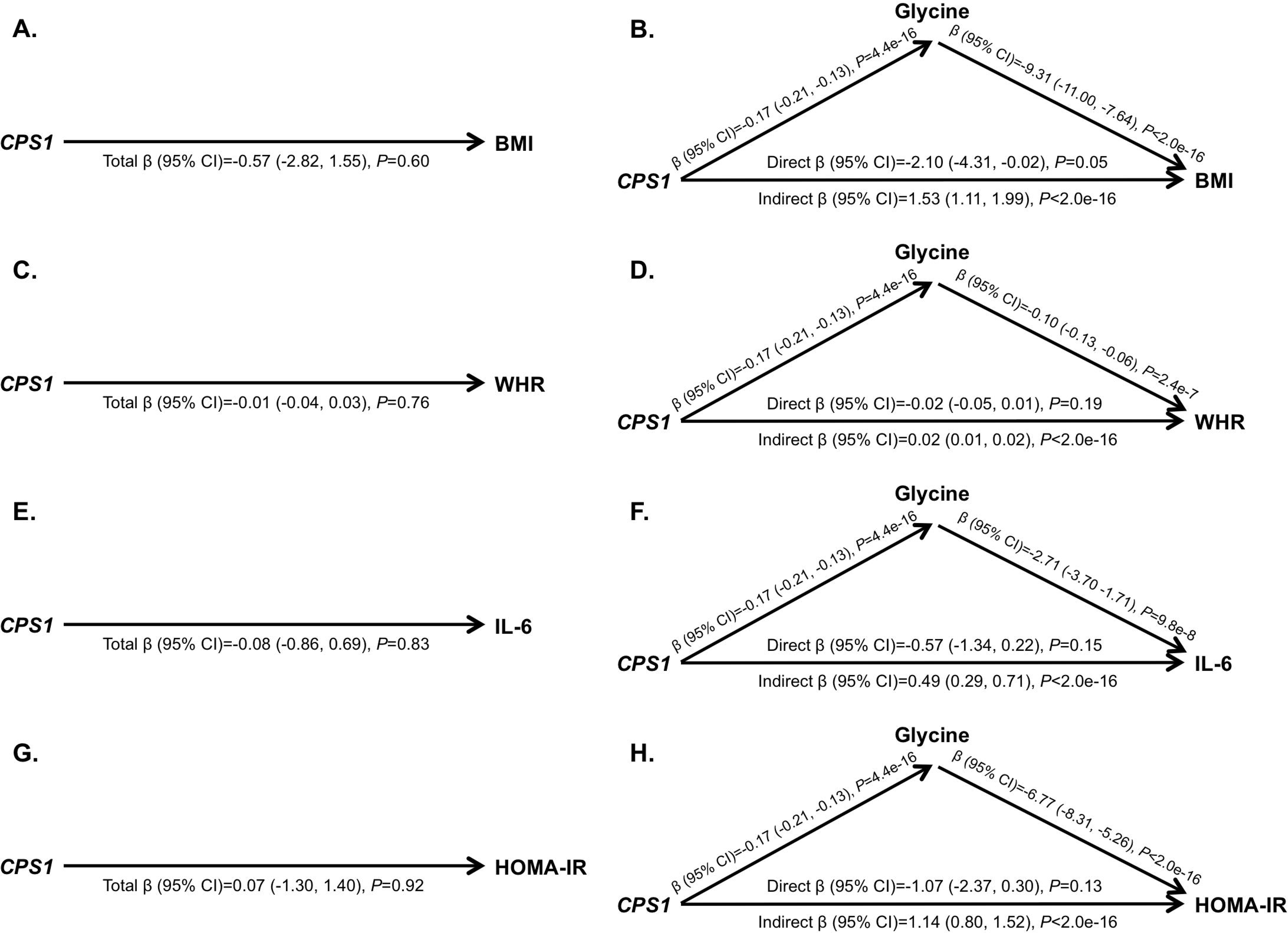
Mediation analyses to assess whether plasma glycine mediates the relationships between imputed *CPS1* expression, BMI, WHR, IL-6, and HOMA-IR. A. Total effect of *CPS1* on BMI. B. Direct and indirect effects of *CPS1* on BMI. C. Total effect of *CPS1* on WHR. D. Direct and indirect effects of *CPS1* on WHR. E. Total effect of *CPS1* on IL-6. F. Direct and indirect effects of *CPS1* on IL-6. F. Total effect of *CPS1* on HOMA-IR. G. Direct and indirect effects of *CPS1* on HOMA-IR. All models adjusted for age and sex; models including *CPS1* additionally adjusted for the first four PCs; models that included glycine additionally adjusted for cholesterol lowering medication use and sample storage time.

## Discussion

We developed an integrative network to investigate relationships between genomics, plasma metabolomics, CSF metabolomics, and AD risk factors. Although no gene expression levels were directly correlated with AD risk factors, there were many instances of genes being indirectly correlated with AD risk factors though metabolites. Building on one such instance, we found that glycine mediated the pathway between *CPS1* expression and cardiovascular and diabetes risk factors. This suggests that our results may have generated many valid hypotheses that warrant further investigation. We also found that correlations between plasma and CSF metabolites ranged widely but typically had low correlations. This could suggest that most plasma metabolites are not representative of certain metabolic changes occurring in the brain, although we cannot rule out the possibility that the low average correlation is, at least partially, due to the time difference between the plasma and CSF sample collection.

The low correlation we observed between plasma and CSF metabolite levels could be related to ~98% of small molecules not being able to pass the blood-brain barrier (BBB)(57). Cholesterol is an example of a lipid metabolite that typically cannot pass the BBB(58), and was not correlated between tissues (r=−0.07). On the other hand, caffeine (a xenobiotic) readily crosses the BBB(59) and it was highly correlated between tissues (r=0.81), as was 5-acetylamino-6-amino-3-methyluracil (r=0.82), which is a caffeine metabolite, and theophylline (r=0.82), which is structurally and pharmacologically similar to caffeine. This could contribute to lipids having the weakest average correlation and xenobiotics having the strongest average correlation between plasma and CSF tissues. However, it is important to note that metabolites within a given pathway can vary widely from each other and should be considered on an individual basis, accordingly, as the averages presented here may not reflect a particular metabolite’s unique properties. The hypothesis about plasma and CSF differing due to the BBB is also supported by the only correlations in the network analysis involving CSF biomarkers (*i.e.*, tau) being with CSF metabolites, although we cannot rule out the possibility that this correlation is related to CSF biomarkers and CSF metabolomics being analyzed from the same sample and thus, not having time-related variation.

Our network analysis revealed that 38 CSF metabolites were highly predictive of CSF T-tau and P-tau, collectively explaining 60.7% and 64.0% of the variance of T-tau and P-tau, respectively. Further investigations of these CSF metabolites could lead to a better understanding of mechanisms and pathways that influence the development of tau tangles. In contrast, no CSF metabolites were correlated with CSF amyloid biomarkers, which could have implications about the biological function of amyloid versus tau. It is possible that we did not capture the small molecules that amyloid may be associated with, or that amyloid is generally not associated with small molecules. Although our CSF findings were limited by their small sample size, they offer potentially novel information regarding the interface between CSF biomarkers and CSF metabolites, as we have not identified previous studies investigating these relationships.

One advantage of using imputed gene expression data is that it only represents the genetically regulated component of gene expression, reducing the risk of confounding due to environmental factors and reverse causality in mediation analyses. We found that glycine mediated the relationship between *CPS1* and BMI, WHR, IL-6, and HOMA-IR, such that lower *CPS1* expression was associated with higher levels of glycine, which were associated with lower BMI, WHR, IL-6, and HOMA-IR. Relationships between *CPS1*, glycine, and cardiovascular risk factors have been hypothesized recently, but not clearly defined(43, 60). The *CPS1* (Carbamoyl-Phosphate Synthase 1) gene encodes for a mitochondrial enzyme that catalyzes the first step of the hepatic urea cycle by synthesizing carbamoyl phosphate from ammonia, bicarbonate, and two molecules of ATP, and is important for removal of urea from cells(61). Notably, all genes encoding enzymes involved in the urea cycle are expressed in the brain, including *CPS1*(62), and levels of enzymes and metabolic intermediates involved in the urea cycle are altered in AD patients(63). *CPS1* variants have been linked to CPS1 deficiency(61), neonatal pulmonary hypertension(64), vascular function(65), traits related to blood clotting, such as fibrinogen levels and platelet count(66–69), homocysteine levels(70–73), HDL cholesterol(74), kidney function and disease(75–78), AD(79), and BMI(80, 81). Higher adipose tissue expression of *CPS1* has been associated with detrimental traits, including weight gain(60). At least nine studies have reported associations between *CPS1* variants and glycine(15, 16, 41–47) and others have reported associations with betaine, a derivative of glycine(15, 16, 82). Glycine is a common amino acid involved in the production of DNA, phospholipids, and collagen, and in the release of energy. Previous studies have identified negative correlations between glycine and cardiovascular and diabetes risk factors such as BMI, waist circumference, HOMA-IR, obesity and visceral obesity, subcutaneous and visceral fat area, hypertension, and acute myocardial infarction(45, 48–55). These previous findings are in the same direction as our findings and are highly supportive of the biological relevance of our results, which lead us to hypothesize that the *CPS1*-cardiovascular risk pathway is linked through the mediation of glycine.

One particular *CPS1* variant, rs715, has been linked to urine and blood glycine levels(15, 16, 43–45), blood levels of betaine(15, 16, 82), blood levels of fibrinogen(66, 67), and BMI(80). This is a common variant, with a MAF=0.27 based on 62,784 whole genome sequences from Trans-Omics for Precision Medicine (TOPMed)(83). The minor C allele of rs715 decreases *CPS1* expression(82). To further test our findings, we conducted additional mediation analyses using this variant and found highly consistent results, suggesting that having one or two minor alleles of rs715 (which decreases *CPS1* expression) increases levels of the three glycine plasma metabolites, which decreases BMI, WHR, IL-6, and HOMA-IR (Figures S9-S11). Thus, the minor C allele of rs715 may have a protective role in cardiovascular risk.

One of the primary strengths of this analysis is that it shows the feasibility of performing integrated omics analyses and the potential utility of such approaches. It is becoming more common for cohorts to collect such datasets; for example, the National Institutes of Health is sponsoring the new TOPMed nation-wide consortium that aims to deeply phenotype its participants utilizing omics technologies (www.nhlbiwgs.org). It is anticipated that initiatives such as TOPMed will greatly advance our knowledge of many complex diseases and traits. However, to fully utilize these rich data, it will be crucial to identify effective means of integrating them and maximize their potential to provide a more holistic understanding of the disease process. While there is still a great need for such methods, our inter-omic network analysis and subsequent targeted follow-up analyses outlines one approach to effectively integrate omics data.

This study was not without limitations. Due to computational burdens, our network analysis did not fully utilize the longitudinal aspect of our data. Further, our sample sizes for CSF biomarkers and metabolites were limited, which is likely why we had few CSF findings in our network analysis. Plasma and CSF samples typically were not collected on the same day, which could influence our correlation results. However, this may not have influenced our network analysis to a large extent because we averaged the residuals of longitudinal traits. We were unable to include smoking behavior in our network analysis due to the prohibitive number of smokers in our cohort (n=48). Despite these limitations, we were encouraged to find that many of our results had been previously reported, thereby strengthening confidence in our novel findings.

Opponents of the “big data” era have criticized omics approaches because they are not hypothesis driven and do not follow the standard scientific method(84, 85). However, we know biology to be complex far beyond our current understanding. To believe that we currently have the ability to generate valid biological hypotheses to understand complex conditions without data would be a fallacy. This was a lesson learned in the years preceding the completion of the human genome sequence in 2001 when research efforts were heavily invested into targeted genetic loci and genome-wide linkage screens of ~500 loci(86). This approach was successful for genes that follow Mendelian patterns, such as highly penetrant variants in the *BRCA1* and *BRCA2* genes that are responsible for inherited forms of breast cancer and the *APP*, *PSEN1*, and *PSEN2* genes that cause the inherited early onset form of AD. However, it had limited success for traits that follow complex inheritance patterns(86). The utility of omics data, and particularly integrated omics approaches, is the ability to generate data driven hypotheses. Our knowledge of biology has been evolving for centuries; however, with the data we are able to generate due to recent biotechnological advances, we now have the opportunity to advance our knowledge of biology at an unprecedented rate. Such data could lead to dramatic improvements in the state of preventative and therapeutic medicine, particularly for complex diseases such as AD, for which few such preventative or therapeutic methods exist and little is known about the underlying biological mechanisms.

By integrating genomics, metabolomics, and clinical risk factors for AD, we were able to identify complex relationships that offer insight into the onset of AD and risk factors associated with its onset. Our research has generated many promising hypotheses that could drive subsequent experimental investigations and potentially offer clinicians and researchers new insights regarding the development of tau tangles. As the generation of omics data accelerates across investigations of a variety of research fields, continued efforts to navigate statistical and computational issues will be critical. The work presented here represents early efforts to integrate omics data, but much more research is needed to identify the most effective means of doing so and thereby maximize the utility of such rich sources of data. The success of precision medicine is heavily reliant on the advancement of computational biology and the ability to translate millions of biological data points into individual clinical implications.

## Acknowledgements

The authors thank the University of Wisconsin Madison Biotechnology Center Gene Expression Center for providing Illumina Infinium genotyping services. We especially thank the WRAP participants.

